# The Enemy Within: how pathogen-free are your commercial specific pathogen-free mice?

**DOI:** 10.1101/2024.07.05.602177

**Authors:** David Mark, Mia Trolan, Gillian Douce, Andrew J Roe, Rebecca E McHugh

## Abstract

Here, we report the incidental isolation and genome sequencing of an *E. coli* O157:H7 strain isolated from the faeces of commercial, specific-pathogen free mice (BALB/c). The isolate, subsequently named EC_MP01 was found to carry both Shiga toxin 2a and the Locus of Enterocyte Effacement (LEE) pathogenicity island, major virulence factors associated with severe disease in humans. This report draws attention to the limitations of existing screening systems and highlights how strain carriage influence the outputs from such studies.

## Main Body

Animal models of bacterial infection are routinely used to study host-pathogen interactions and test novel antimicrobial therapies. Therefore, commercial mice used in these experiments must be free of pathogens which may affect the outcome or reproducibility of experiments. Specific Pathogen Free (SPF) mice are bred for this purpose, and they are routinely screened for microorganisms which are known to cause disease in mice^1^. These include bacterial pathogens such as Beta-haemolytic *Streptococcus, Citrobacter rodentium* and *Salmonella spp*, and >40 additional bacterial, viral and fungal pathogens. Culture-based methods including serology are typically used in screening, and in more recent years, PCR-based methods have been implemented. Notably, *Escherichia coli* is not routinely screened for by many commercial breeders as human intestinal *E. coli* pathotypes do not readily cause enteric disease in mice and therefore are unlikely to compromise experimental outcomes.

The inability for these *E. coli* strains to cause disease in mice has necessitated the development of a specific murine model of enteropathogenic *E. coli-*mediated disease. *Citrobacter rodentium* is a natural murine pathogen which is a causative agent of transmissive colonic hyperplasia in laboratory mice^2^. The virulence factors used by *C. rodentium* used to colonize the murine gut mirror those found in Enteropathogenic and Enterohaemorrhagic *E*.*coli* (EPEC/EHEC), and therefore it is the archetypal model of attaching and effacing bacterial pathogenesis in mice^3^. In our experiments, mice are infected with *Citrobacter rodentium* DBS770, a Shiga-toxin encoding derivative designed to mimic human STEC pathology in mice^4^. This allows for the testing of novel therapeutic disease in a model of STEC, for which there are currently no antibiotic therapies available^5^.

The central aim of our planned experiment was to evaluate the efficacy of the anti-virulence drug aurodox against *C. rodentium* DBS770 infections in mice. In addition, DNA was extracted from the faeces of mice at five points throughout the fourteen day experiment for microbiome analysis via 16S amplicon sequencing. A total of four groups of six animals were used, two groups were mock-infected, with one cage treated with vehicle only, and the other treated with aurodox. The two remaining groups of animals were infected with *Citrobacter rodentium* + Stx DBS770, with one cage was treated with vehicle alone and the other with aurodox. As our aim from the outset of the experiment was to establish how aurodox treatment may influence the microbiome, DNA was extracted from faeces collected on Day 0 (before the procedure began), Day 3, Day 7, Day 10 and Day 14 of the experiment for amplicon sequencing of the faecal microbiome. In addition, colony-forming unit counts on selective chloramphenicol agar were carried out for infected animals on alternative days, starting on Day 1. Early in the experiment, it was observed that the colonies isolated from the infected group of mice treated with aurodox were heterogeneous when growing on agar containing chloramphenicol, with some colonies appearing larger in diameter (Figure 1A). On day 3, faecal samples from uninfected animals were screened for cross-cage contamination using culture on selective agar (chloramphenicol). This process lead to the isolation of mucoid, chloramphenicol-resistant colonies from the faeces of animals which had not been infected with *Citrobacter rodentium* during the experiment (Figure 1B), but had been treated with aurodox. To identify the species of the contaminating organism, 16S rRNA sequencing was performed (Figure 1C). The organism was identified as *Escherichia coli*.

**Figure 1:**
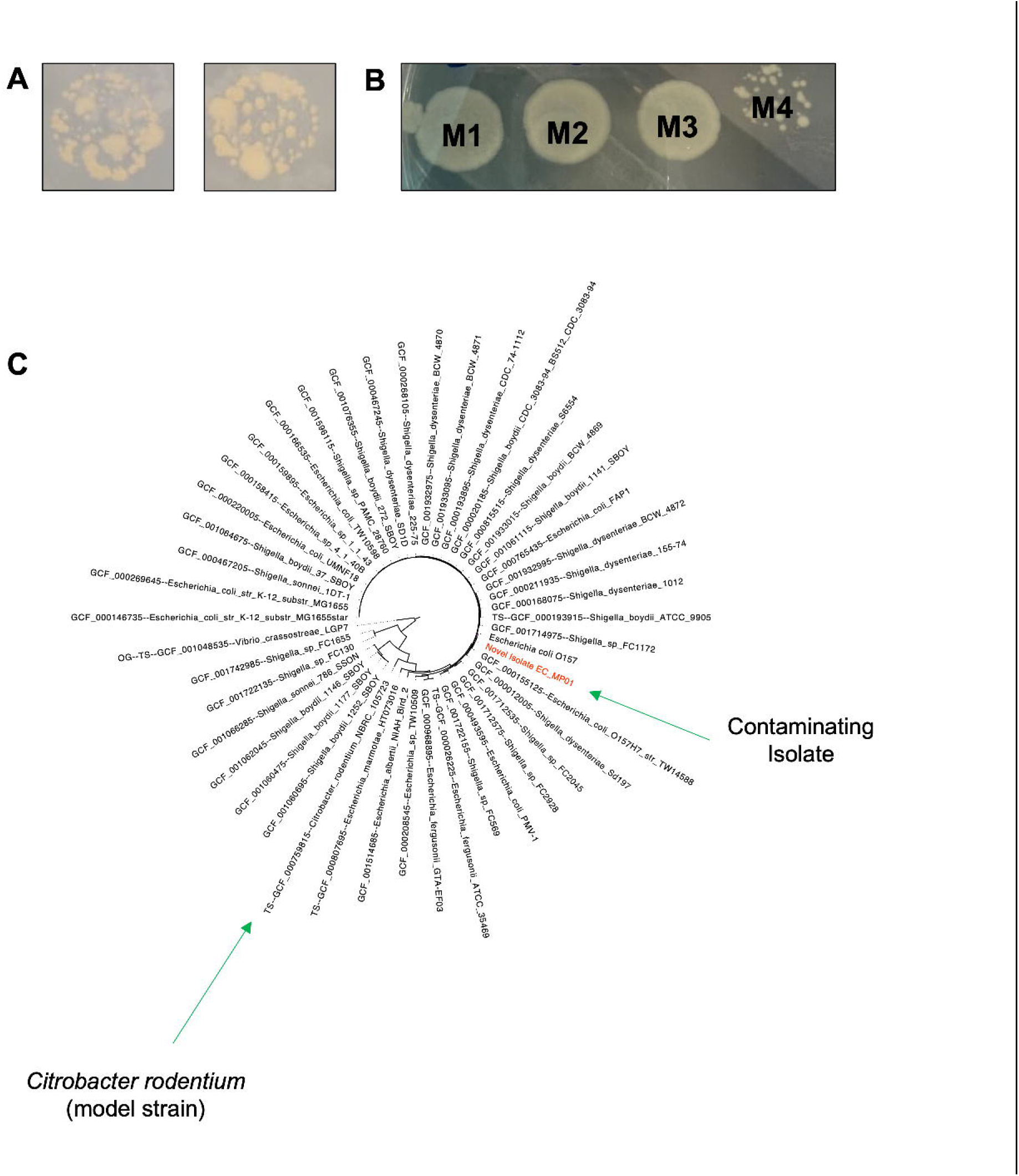
Identification of contaminating colonies on selective chloramphenicol agar. (A) Observation of heterologous morphology between colonies. Photographs show bacteria isolated from murine faeces from BALB/c mice infected with *Citrobacter rodentium* D8S770, treated with corn oil vehicle (left) and 25 mg/kg aurodox (right). (B) Isolation of mucoid, chloramphenicol resistant colonies from uninfected mice. Bacterial isolation from four mice (M1-M4) is shown. (C) Phylogenetic tree depicting phylogeny of isolate EC_MP01. AutoMLST (Alanjary, Steinke, & Ziemert, 2019) was used to build phylogenetic tree based on MASH distance value. FigTree ® software was used to format the tree. Position of EC_MP01 is highlighted in red.

Due to the high similarity of 16S rRNA nucleotide sequence between *Escherichia coli* and *Citrobacter rodentium* (>97%), further analysis was required to confirm that the species discovered in the faeces of uninfected mice was truly *E. coli*, and not *C. rodentium* DBS770 potentially introduced to the mice through cross-contamination. Therefore, whole genome sequencing of the isolate, subsequently named EC _MP01, was performed. To maximise the quality of the sequence, a combination of Illumina and Oxford Nanopore Sequencing was used. This resulted in the generation of a high-quality genome assembly (N50= 5,291,832, L50=1, Contig number=32, Figure S1). Analysis of this assembly using QUAST^6^ and visualisation using Bandage^7^ revealed the presence of one large 5.3 Mb chromosome and multiple mobile genetic elements including a 90 kb plasmid and 64 kb phage. This assembly can be found in full on NCBI under Bioproject PRJNA1131981.

To confirm that the isolate was *E. coli* and assign taxonomic labels to each of the contigs, Kraken analysis was undertaken^8^. This revealed that all contigs were assigned to the genus *Escherichia*, with *Escherichia coli* representing the most frequently identified species. Subsequently, AutoMLST^9^ was used to determine the most closely related strains to EC_MP01. Alarmingly, the closest related genome in the database was *E. coli* EDL933 with an average nucleotide identity of 99.9% and a MASH distance of 0.0015. A whole genome BLAST between EC_MP01 and EDL933 was executed via the Proksee server^10^ (Figure S1) which confirmed that the EDL933 chromosome and the archetypal pO157 plasmid comprise the core of the EC_MP01 genome.

To gauge the potential risk of this pathogen to personnel handling these animals, analysis of virulence genes within EC_MP01 was carried out. This identified genes encoding Shiga Toxin 2a, which is the toxin most associated with the development Haemolytic Uremic Syndrome (HUS) in humans^11^. Furthermore, the strain encodes a complete Locus of Enterocyte Effacement (LEE) pathogenicity island, with >99% similarity to the LEE of EDL933 (Figure 2C). In STEC pathogens, this pathogenicity island facilitates the attachment and effacement to the epithelial cells of the gut and therefore carrying this demonstrates the ability of the pathogen to cause disease in humans. In addition, pLannotate^12^ was used to identify two additional plasmids (3.5 kb and 5.5kb) in the EC_MP01 genome which were absent in EDL933. Remarkably, one of the plasmids, subsequently named pStx_GFP, was found to encode a transcriptional-fusion GFP reporter system controlled under the Stx_2a promoter. Sequence depth for this plasmid was 23 x higher than the chromosome confirming that multiple copies exist within each cell.

**Figure 2:**
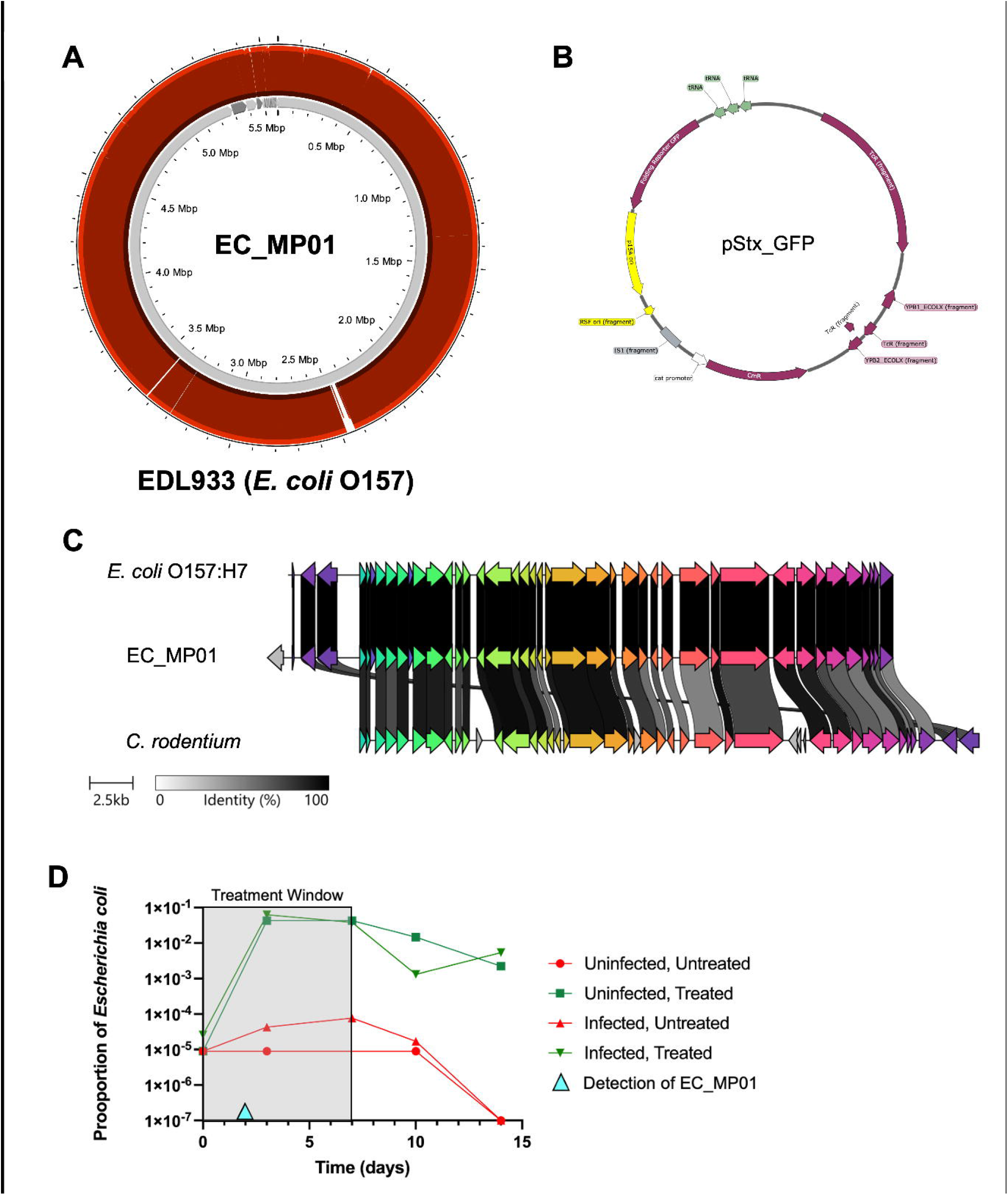
Analysis of the EC_MP01 genome. (A) Mapping of the EC_MP01 genome against the genome of the most closely related strain in the database, EDL933. Analysis carried out via BLAST function in Proksee. Inner track represents EC_MP01, outer track represents EDL933. (B) Map of the 5.5 kb plasmid pStx_GFP, encoded by EC_MP01. Map generated by Snapgene^™^. (C) Alignment of LEE pathogenicity islands of *E*.*coli* O157:H? (EDL933), EC_MP01 and C. *rodentium* D8S100. Figure generated by Clinker^™^.

**Figure 4:**
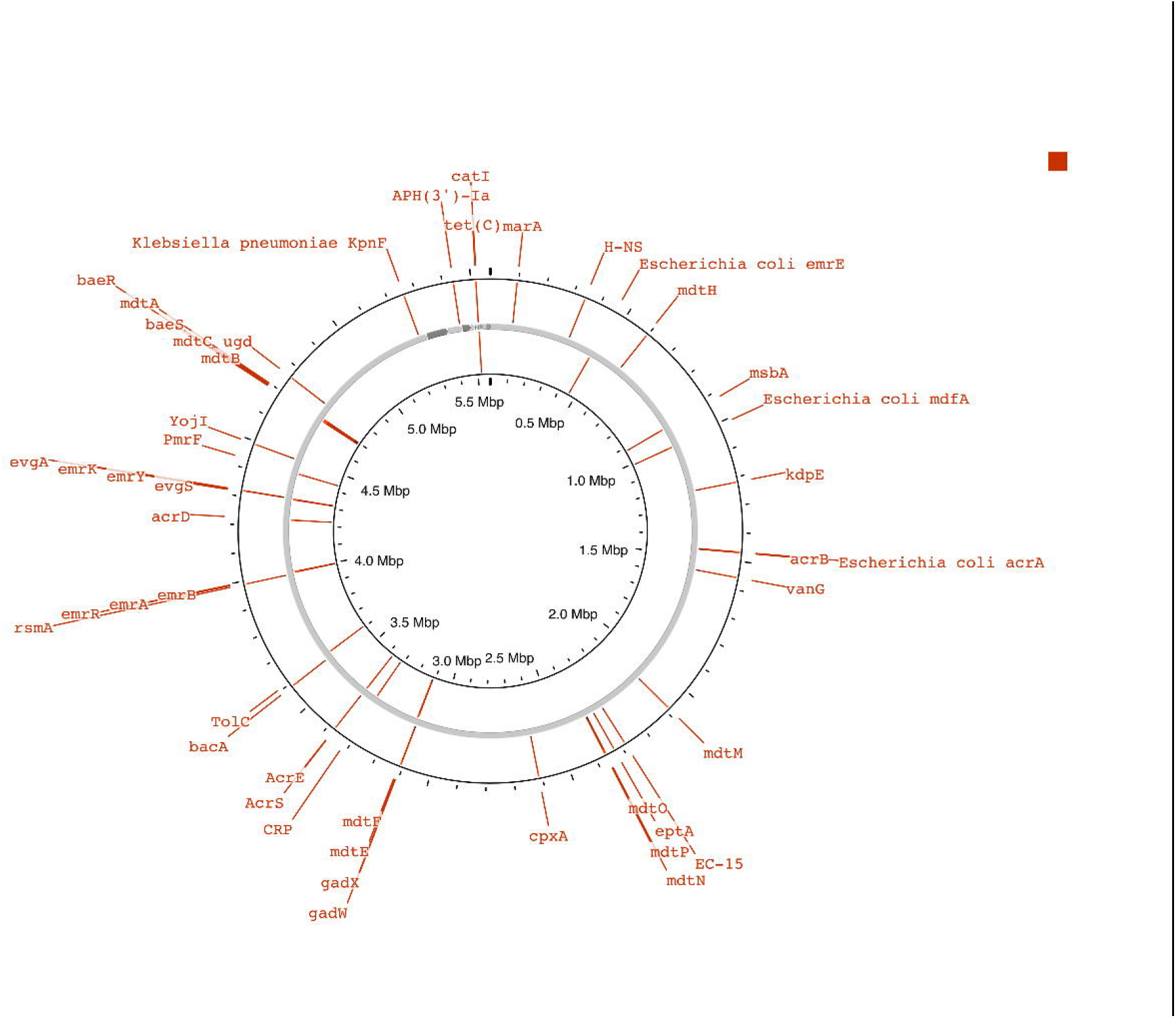
Analysis of Antimicrobial Resistance genes encoded by EC_MP01. AMR-associated genes were identified using CARD (comprehensive antibiotic resistance database) function of Proksee. A full list of targets, mutations and classes of antibiotics can be found in Supplementary Figure S3.

One feature of ECMP_01 which initiated this investigation, was the ability of the strain to grow on selective agar containing chloramphenicol. To investigate the chloramphenicol resistance phenotype of ECMP_01 and gauge an understanding of its resistance/sensitivity to other antibiotics, CARD analysis of the ECMP_01 genome was performed (Figure S2). Here, multiple chloramphenicol resistance genes were identified including a classic acyltransferase on the pStx_GFP reporter plasmid. In addition, a further 55 hits for AMR genes (57 in total) were identified which have the potential to encode resistance to tetracyclines, kanamycin, streptomycin and beta-lactamases (Table S1).

Overall, this work identified the carriage of a Shiga toxin-producing strain of *E. coli*, that appeared to be multi-antibiotic resistant and which carried an engineered fluorescent reporter plasmid. This classifies this isolate as a UK Hazzard Level 3 pathogen, which is not used within our institute. We believe that recovery of the organism in these experiments was the result of the ‘perfect experimental storm’. It has previously been shown that aurodox inhibits *E. coli* O157 epithelial cell attachment^13^, and interestingly shedding of EC_MP01 was only observed in mice treated with the compound. This compounded with use of chloramphenicol selection, to which the strain was resistant allowed the isolation of a strain which may have been previously undetected in the faecal bacteriome of this SPF-mice. This hypothesis is supported by 16S amplicon sequencing which highlights an increased abundance of *E. coli* in the faeces of aurodox-treated mice (Figure 2D). This observation does reduce the concern for personnel handling animals in which this strain could form a minor, almost undetectable component of the microbiome. However, it does have implications for those studying virulence inhibitors *in vivo* as consideration of the potential to isolate pathogens co-incidentally should be considered. There are also potential consequences for experimental outcomes. For example, mice with low levels of EC_MP01 in the gut may have developed immunity to the virulence mechanisms of *E. coli* O157, which are mirrored in *C. rodentium*. In summary, the isolation of this strain highlights that the faecal bacteriome of commercial mice remains poorly characterised, and screening should include metagenomic methods to rule out the presence of pathogens at low levels which may implicate experimental outcomes or compromise safety.

## Acknowledgements

For the purpose of open access, the author(s) has applied a Creative Commons Attribution (CC BY) licence to any Author Accepted Manuscript version arising from this submission. We would like to acknowledge Dr Alice Gallagher, University of Glasgow Biosafety Advisor for her advice.

## Animal Study

All procedures were performed in strict accordance with the Animals (Scientific Procedures) Act 1986 with specific approval granted by the Home Office, UK (PPL PI440270). These applications were also considered and approved by the University of Glasgow Animal Welfare Ethical Review Body (AWERB). Food and water were provided *ad libitum* and animals kept at a constant room temperature of 20–22 °C with a 12 h light/dark cycle.

To prepare the inoculum for oral challenge, an overnight culture of *Citrobacter rodentium* DBS770^1^ grown in 5 ml of LB broth was pelleted via centrifugation and resuspended in PBS to a concentration of 5 × 10^9^ CFU/ml. Two groups of four Specific Pathogen Free, Balb/C mice were either orally challenged with 1 × 10^9^ CFU *C. rodentium* DBS770 in a 200 µl (infected) volume, or an equivalent amount of PBS (uninfected). Due to the commercial sensitivities associated with this publication, the commercial breeder will not be named. For the two groups of animals infected with *C. rodentium*, one was treated with 200 µl corn oil containing 25 mg/kg aurodox and the other with 200 µl corn oil alone (vehicle-treated). Two groups of uninfected control animals were similarly treated. Animals were monitored daily for appearance of clinical symptoms and/or weight loss and were given a soft food diet if weight loss >10%. Animals which lost 20% of their body weight were culled. Faecal samples were collected for CFU counts on alternate days throughout the experiments, beginning on day 1 and on Day’s 0, Day 3, Day 7, Day 10 and Day 14 for 16S amplicon metagenomics analysis.

### 16S Amplicon Sequencing of EC_MP01

The 16S rRNA gene from EC_MP01 was amplified from faecal isolates from uninfected mice (Day 2), GoTaq Green standard colony PCR conditions with forward primer AGAGTTTGATCCTGGCTCAG and reverse primer GGTTACCTTGTTACGACTT^2^ were used. Amplicons were Sanger sequenced (Eurofins, UK) and BLASTn of resulting sequence was carried out to determine bacterial species.

### 16S Amplicon Metagenomic Sequencing

DNA was extracted from fresh faecal samples using the DNeasy® PowerSoil ® Pro Kit (Qiagen). These samples were sent to Novogene (Cambridge, UK) for 16S metagenomic amplicon sequencing. Briefly, library preparation was carried out via amplification of the V3-V4 region of the 16S rRNA gene, (forward primer: CCTAYGGGRBGCASCAG, reverse primer GGACTACNNGGGTATCTAAT). Sequencing reads were acquired using the Illumina Novaseq 6000. Bioinformatic analysis and abundance of species was carried out in house,

### Whole Genome Sequencing and analysis

Sequencing was carried out by Microbes NG, Birmingam UK, according to their protocols. For Illumina sequencing, genomic DNA libraries were prepared using the Nextera XT Library Prep Kit (Illumina, San Diego, USA) following the manufacturer’s protocol. Genome Sequencing Methods DNA quantification and library preparation were performed on a Hamilton Microlab STAR automated liquid handling system (Hamilton Bonaduz AG, Switzerland). Libraries were sequenced on an Illumina NovaSeq 6000 (Illumina, San Diego, USA) using a 250 bp paired-end protocol. Reads were adapter trimmed using Trimmomatic version 0.30^3^ with a sliding window quality cut-off of Q15. For Oxford Nanopore sequencing, long-read genomic DNA libraries are prepared with Oxford Nanopore SQK-LSK109 kit with Native Barcoding EXP-NBD104/114 (ONT, United Kingdom) using 400-500 ng of HMW DNA. Barcoded samples were pooled together into a single sequencing library and loaded in a FLO-MIN106 (R.9.4.1) or FLO-MIN111 (R10.3) flow cell in a GridION (ONT, United Kingdom). Hybrid assembly is performed using Unicycler version 0.4.0, and contigs were annotated using Prokka^4^.

### Safety Statement

Upon initial indication of Shiga toxin carriage, all biological material harvested from this experiment including EC_MP01 was immediately destroyed by autoclaving and disposed of according to biological risk assessments under guidance from the Biological Safety Officer (University of Glasgow).

---

**Figure S1:**
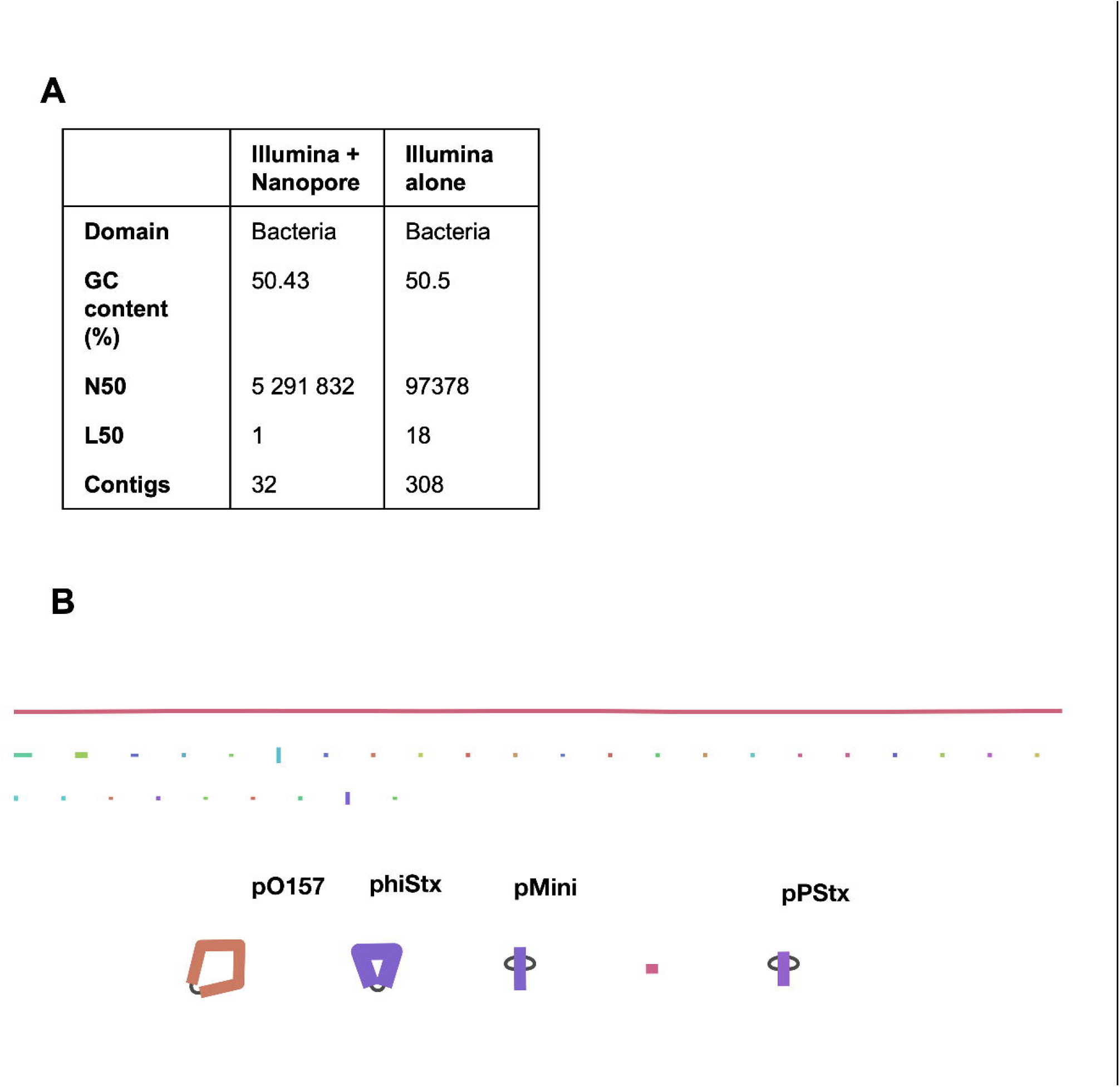
Quality Analysis of EC_MP01 genome. (A) Quast analysis of EC_MP01 genome assemblies for lllumina sequencing alone vs lllumina and Oxford Nanopore. (B) Bandage depiction of EC_MP01 hybrid assembly. Linear chromosome (5.3 Mb) is depicted as red line with smaller contigs represented in the coloured lines underneath. Selected mobile genetic elements including p0157, phiStx and pStx are highlighted.

**Table S3:**
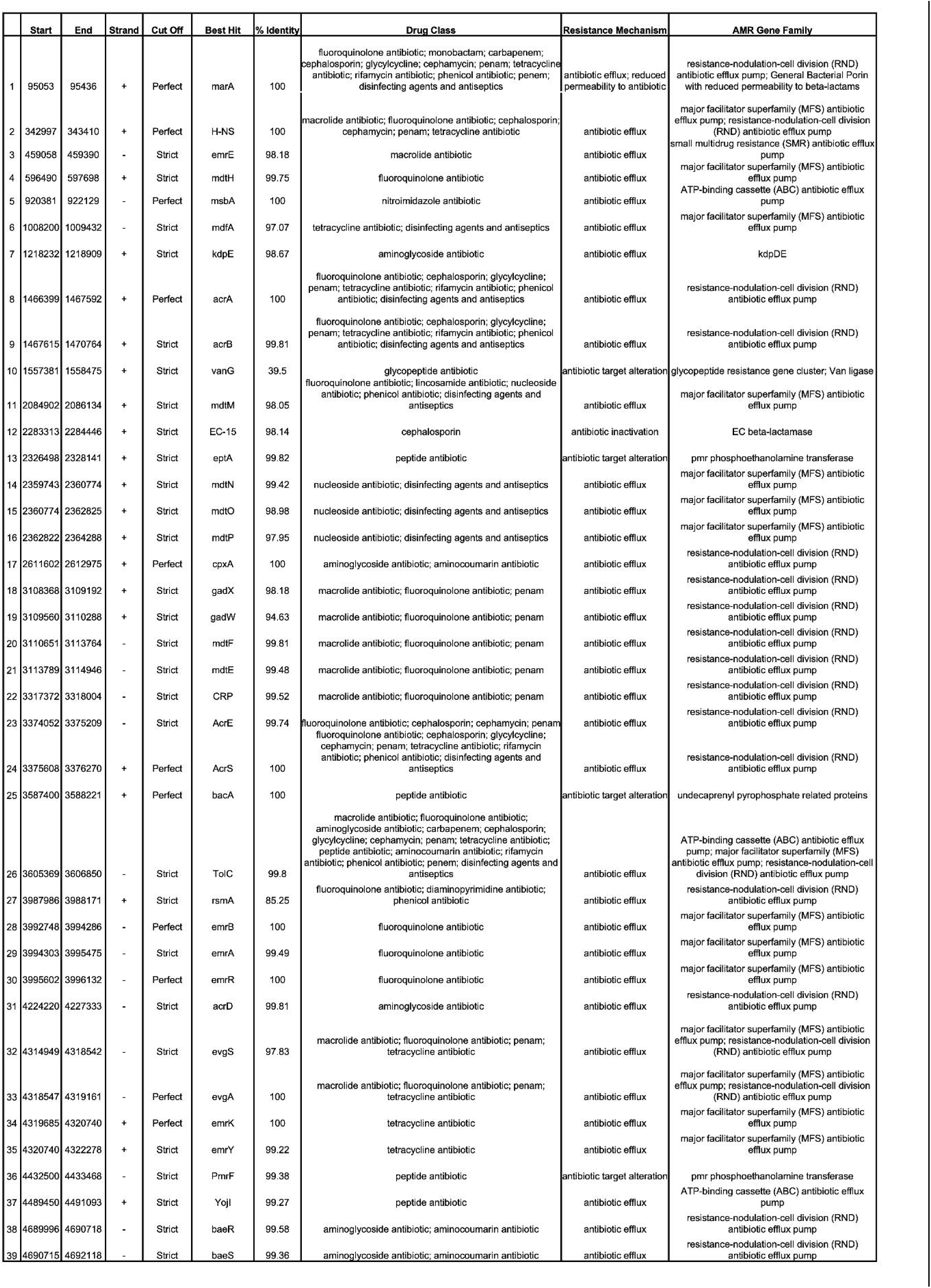
Detailed list of AMR-associated genes as detected by CARD.

## Bibliography

1 Foster, H. L. The development of specific pathogen free and germfree animals. Biomed Purv 1, 76–85 (1961).

2 Crepin, V. F., Collins, J. W., Habibzay, M. & Frankel, G. Citrobacter rodentium mouse model of bacterial infection. Nature Protocols (2016). 10.1038/nprot.2016.100

3 Deng, W. et al. Dissecting virulence: Systematic and functional analyses of a pathogenicity island. Proceedings of the National Academy of Sciences of the United States of America 101, 3597–3602 (2004). 10.1073/pnas.0400326101

4 Mallick, E. M. et al. A novel murine infection model for Shiga toxin-producing Escherichia coli. Journal of Clinical Investigation 122, 4012–4024 (2012). 10.1172/JCI62746

5 Tarr, P. I. & Freedman, S. B. Why antibiotics should not be used to treat shiga toxin-producing escherichia coli infections. Current Opinion in Gastroenterology 38, 30–38 (2022). 10.1097/MOG.0000000000000798

6 Gurevich, A., Saveliev, V., Vyahhi, N. & Tesler, G. QUAST: Quality assessment tool for genome assemblies. Bioinformatics (2013). 10.1093/bioinformatics/btt086

7 Wick, R. R., Schultz, M. B., Zobel, J. & Holt, K. E. Bandage: interactive visualization of de novo genome assemblies. Bioinformatics 31, 3350–3352 (2015). 10.1093/bioinformatics/btv383

8 Davis, M. P., van Dongen, S., Abreu-Goodger, C., Bartonicek, N. & Enright, A. J. Kraken: a set of tools for quality control and analysis of high-throughput sequence data. Methods 63, 41–49 (2013). 10.1016/j.ymeth.2013.06.027

9 Alanjary, M., Steinke, K. & Ziemert, N. AutoMLST: an automated web server for generating multi-locus species trees highlighting natural product potential. Nucleic acids research 47, W276–W282 (2019). 10.1093/nar/gkz282

10 Grant, J. R. et al. Proksee: in-depth characterization and visualization of bacterial genomes. Nucleic Acids Res 51, W484–W492 (2023). 10.1093/nar/gkad326

11 Melton-Celsa, A. R. Shiga Toxin (Stx) Classification, Structure, and Function. Microbiology Spectrum (2014). 10.1128/microbiolspec.ehec-0024-2013

12 McGuffie, M. J. & Barrick, J. E. pLannotate: engineered plasmid annotation. Nucleic Acids Res 49, W516–W522 (2021). 10.1093/nar/gkab374

13 McHugh, R. E., O’Boyle, N., Connolly, J. P. R., Hoskisson, P. A. & Roe, A. J. Characterization of the mode of action of aurodox, a type III secretion system inhibitor from streptomyces goldiniensis. Infection and Immunity (2019). 10.1128/IAI.00595-18

## References

1 Mallick, E. M. et al. A novel murine infection model for Shiga toxin-producing Escherichia coli. Journal of Clinical Investigation 122, 4012–4024 (2012). 10.1172/JCI62746

2 Frank, J. A. et al. Critical evaluation of two primers commonly used for amplification of bacterial 16S rRNA genes. Appl Environ Microbiol 74, 2461–2470 (2008). 10.1128/AEM.02272-07

3 Bolger, A. M., Lohse, M. & Usadel, B. Trimmomatic: A flexible trimmer for Illumina sequence data. Bioinformatics (2014). 10.1093/bioinformatics/btu170

4 Seemann, T. Prokka: Rapid prokaryotic genome annotation. Bioinformatics (2014). 10.1093/bioinformatics/btu153

